# Pooled optical screens in human cells

**DOI:** 10.1101/383943

**Authors:** David Feldman, Avtar Singh, Jonathan L. Schmid-Burgk, Anja Mezger, Anthony J. Garrity, Rebecca J. Carlson, Feng Zhang, Paul C. Blainey

**Affiliations:** Broad Institute of MIT and Harvard, Cambridge, Massachusetts, USA; Department of Physics, MIT, Cambridge, Massachusetts, USA; Department of Genetics, Stanford University, Stanford, California, USA; Department of Medical Biochemistry and Biophysics, Karolinska Institute, Stockholm, Sweden; Department of Health Sciences and Technology, MIT, Cambridge, Massachusetts, USA; Department of Biological Engineering, MIT, Cambridge, Massachusetts, USA; McGovern Institute for Brain Research at MIT, Cambridge, Massachusetts, USA; Department of Brain and Cognitive Science, MIT, Cambridge, Massachusetts, USA; Howard Hughes Medical Institute, MIT, Cambridge, Massachusetts, USA

## Abstract

Large-scale genetic screens play a key role in the systematic discovery of genes underlying cellular phenotypes. Pooling of genetic perturbations greatly increases screening throughput, but has so far been limited to screens of enrichments defined by cell fitness and flow cytometry, or to comparatively low-throughput single cell gene expression profiles. Although microscopy is a rich source of spatial and temporal information about mammalian cells, high-content imaging screens have been restricted to much less efficient arrayed formats. Here, we introduce an optical method to link perturbations and their phenotypic outcomes at the singlecell level in a pooled setting. Barcoded perturbations are read out by targeted *in situ* sequencing following image-based phenotyping. We apply this technology to screen a focused set of 952 genes across >3 million cells for involvement in NF-κB activation by imaging the translocation of RelA (p65) to the nucleus, recovering 20 known pathway components and 3 novel candidate positive regulators of IL-1β and TNFα-stimulated immune responses.

## Introduction

Forward genetic screens are a powerful tool for finding genes that cause phenotypes. A variety of methods exist to disrupt genes, introduce exogenous genes, and modulate gene expression in screens of mammalian cells. Many such perturbations can be pooled together and efficiently quantified by next-generation sequencing (NGS) of cell populations. The phenotypic effect of a perturbation can then be defined as an enrichment in cells carrying the perturbation under different conditions (1, 2).

Examples of enrichment-based phenotypes that are compatible with pooled screens include differential cell fitness (e.g., under drug selection) and differential fluorescence of a marker assessed by fluorescence-activated cell sorting (FACS) (e.g. a genetic reporter or immunostained protein) (3). Although such screens have led to important biological discoveries, many complex phenotypes cannot be physically enriched at scale to provide a sample for NGS analysis. As a result, screens of more complex phenotypes have historically been limited to expensive and laborious testing of individual perturbations in arrayed formats. Recently, pooled CRISPR perturbations were integrated with single-cell expression profiling (4–7) to enable screens with a high-dimensional readout of cell state. Alternatively, image-based screens have been used to access a rich set of spatially and temporally defined phenotypes across diverse processes in eukaryotic cells, including mitosis (8, 9), endocytosis (10), viral infection (11), differentiation (12), metabolism (13), DNA damage (14), autophagy (15) and synaptogenesis (16), but have been restricted to arrayed formats. Pooled screens of image-based phenotypes have thus far only been demonstrated in bacterial cells (17, 18).

To address this challenge, we developed a pooled genetic screening protocol compatible with a wide range of spatially and temporally resolved optical assays in mammalian cells, greatly expanding the variety of phenotypes amenable to high-throughput forward genetics. Our approach is to determine both the phenotype and the perturbation identity in each cell by microscopy (**Fig. 1A**), where the perturbations are identified by targeted *in situ* sequencing (19) of associated RNA barcodes. This strategy uses enzymatic amplification and sequencing-by-synthesis (**Fig. 1B**) to provide high signal levels from a compact, easily synthesized barcode. Our method adapts the existing molecular biology pipeline for pooled genetic screens, permitting information-rich phenotyping of thousands of perturbations across millions of cells in a single sample.

**Fig. 1:**
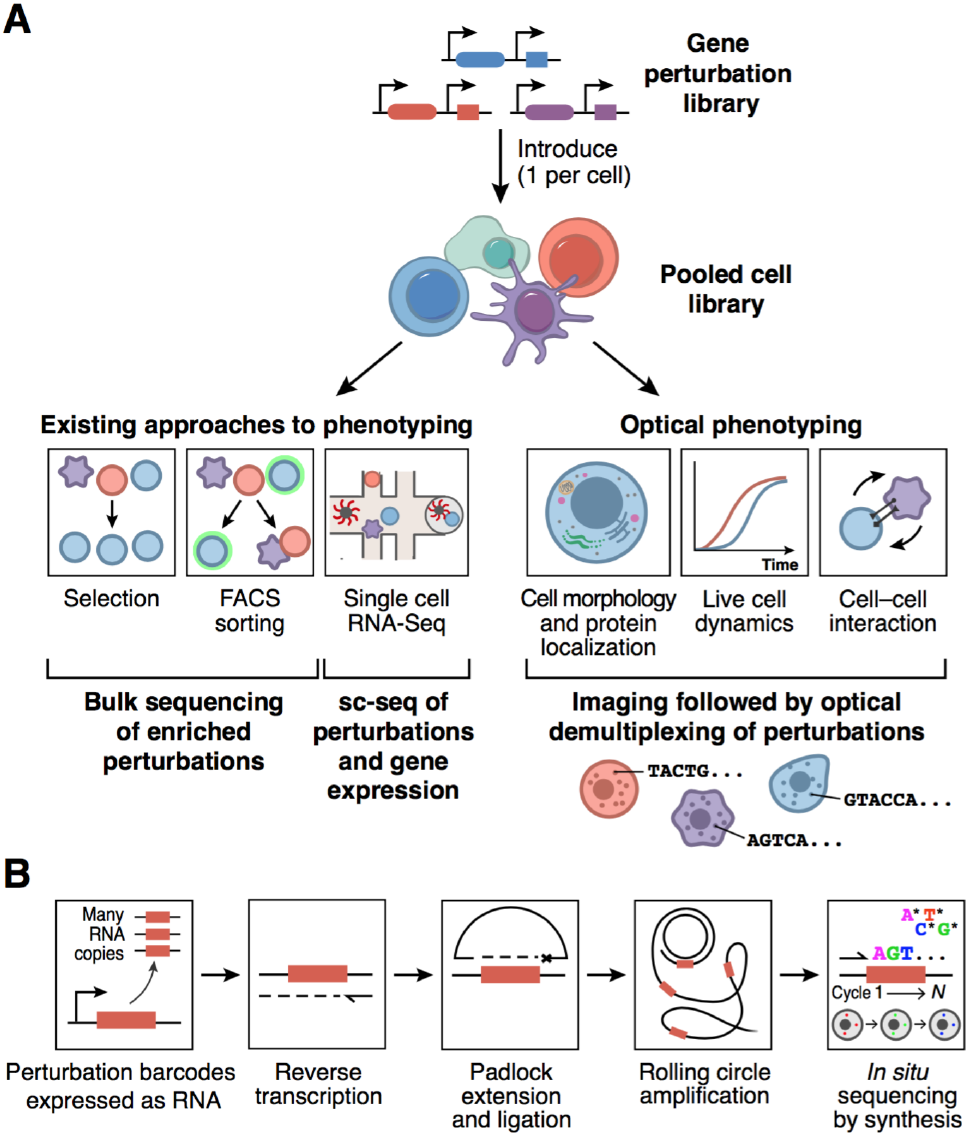
Pooled optical genetic screens. **(A)** In pooled screens, a library of genetic perturbations is introduced, typically at a single copy per target cell. In existing approaches, cellular phenotypes are evaluated by bulk NGS of enriched cell populations or single-cell gene expression profiling. In pooled optical screens, high-content imaging assays are used to extract rich spatiotemporal information from the sample prior to enzymatic amplification and *in situ* detection of RNA barcodes, enabling correlative measurement of phenotype and genotype. **(B)** Targeted *in situ* sequencing is used to read out RNA barcodes expressed from a single genomic integration. Barcode transcripts are fixed in place, reverse transcribed, and hybridized with single-stranded DNA padlock probes, which bind to common sequences flanking the barcode. The 3’ arm of the padlock is extended and ligated, copying the barcode into a circularized ssDNA molecule, which then undergoes rolling circle amplification. The barcode sequence is then read out by multiple rounds of *in situ* sequencing-by-synthesis.

## Results

To apply this approach in the context of CRISPR screening, we encoded the identity of genetic perturbations by cloning guide RNAs (sgRNAs) and associated 12-nt barcodes into lentiGuide-Puro (20) (Addgene #52963), a widely-used sgRNA expression vector (**Fig. 2A, Methods**). Barcodes and flanking sequences were inserted into the 3’ UTR of a Pol II-driven antibiotic resistance gene, a highly expressed mRNA suitable for *in situ* detection, in a design we termed lentiGuide-BC. We used a pooled cloning approach where sgRNAs and barcodes are synthesized in tandem on an oligo array, such that sgRNA-barcode pairings are pre-determined (**Fig. 2A**). As reported for other lentiviral libraries using paired sequences, we initially observed swapping of barcodes and associated sgRNAs in lentiGuide-BC cells due to reverse transcription-mediated recombination (21–24). To address recombination, we utilized a modified lentiviral packaging protocol that reduces barcode swapping from >28% to <5% (25).

To test *in situ* identification of perturbations, we transduced a library of 40 sgRNA-barcode pairs into HeLa-TetR-Cas9 cells at low multiplicity of infection (MOI). We designed a reverse transcription primer and padlock probe complementary to constant regions flanking the barcode, and amplified the barcode via *in situ* reverse transcription, padlock extension/ligation, and rolling circle amplification (**Fig. 1B, Fig. S1 and Table S1**). Sequencing *in situ* using a 4-color sequencing-by-synthesis chemistry yielded bright, compact fluorescent spots that retained base specificity over 12 cycles (**Fig. 2, B and C**). Image segmentation and base calling analysis produced sequence reads with a mapping rate of >85% to the set of known barcodes (>85%) (**Fig. 2D**).

**Fig. 2:**
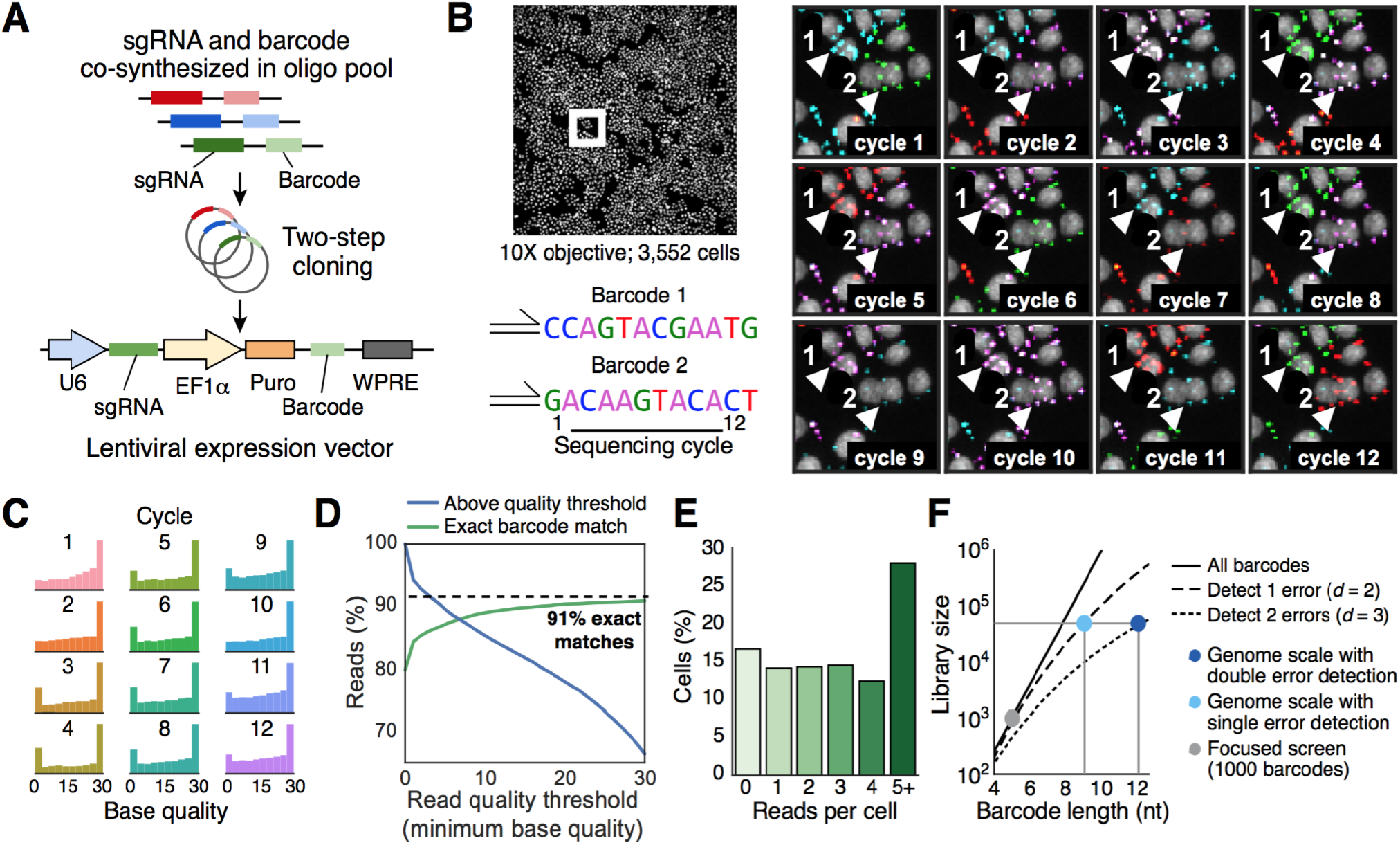
Identification of perturbation barcodes by *in situ* sequencing. (**A**) A 125-nt oligo pool encoding perturbations (sgRNAs) and associated 12-nt barcodes was cloned into a lentiviral vector via two rounds of Golden Gate assembly. (**B**) Expressed barcode sequences were read out by padlock detection, rolling circle amplification, and 12 cycles of sequencing-by-synthesis (data shown for lentiGuide-BC). A linear filter (Laplacian-of-Gaussian, kernel width σ = 1 pixel) was applied to sequencing channels to enhance spot-like features. (**C**) Per-base quality score over 12 cycles of *in situ* sequencing, calculated from signal for called base divided by signal for all bases. (**D**) >80% of barcodes map to 40 designed sequences out of 16.7 million possible 12-nt sequences. (**E**) Most cells contain multiple barcode reads. (**F**) The number of possible barcodes scales geometrically with barcode length. Sufficient 12-nt barcodes can be designed to cover a genome-scale perturbation library while maintaining the ability to detect and reject single or double sequencing errors (minimum pairwise Levenshtein distance d = 2 or 3, respectively).

For optical screening to be scalable, the *in situ* readout step should be able to process millions of cells within a few days, ensuring high coverage per perturbation (typically 100-1,000 cells). This cellular throughput is possible in our system because of the high fluorescence signal-to-noise achieved by padlock-based barcode amplification and sequencing-by-synthesis. High signal enables accurate sequence data to be obtained at low optical magnification in large fields of view, each containing thousands of cells (**Fig. 2B, Movie S1, Tables S2 and S3**). Specifically, following optimization of the barcode amplification protocol including the post-fixation and padlock extension/ligation conditions (**Fig. S2**), *in situ* sequencing signals were readily visible a t 10X optical magnification, with one or more exactly mapped reads detected in over 82% of cells transduced with lentiGuide-BC (**Fig. 2, B and E**).

Following the initial robust detection of 12-nt barcodes, we designed a set of 83,314 barcodes that were used in subsequent screens. As there are 16.7 million possible 12-nt sequences, we selected barcodes with minimum pairwise edit distance of 3, which allowed us to detect and reject up to two errors (insertion/deletion/substitution) arising from oligo synthesis or *in situ* processing (**Fig. 2F, Methods**) (26).

We next sought to evaluate the overall accuracy of mapping perturbation genotype to cell phenotype *in situ.* In principle, errors could arise from oligo synthesis, library cloning, lentiviral delivery, barcode diffusion during *in situ* processing, barcode readout by *in situ* sequencing, or incorrect assignment of reads to cells during image processing. To assess the impact of these error sources on CRISPR-Cas9 screening, we built a lentiviral reporter that produces an HA-tagged, nuclear-localized H2B protein after a Cas9-induced +1 frameshift in a target region (**Fig. 3A, Methods**). Cells expressing the reporter can either be screened *in situ* or by FACS. The reporter is highly specific and sensitive, with a mean *in situ* activation across 5 targeting sgRNAs of 65 ± 2.7% and background of <0.001% in the absence of a targeting sgRNA (**Fig. 3, B and C, Fig. S3**). We transduced cells stably expressing the frameshift reporter with a library containing 972 barcodes redundantly encoding 5 targeting and 5 control sgRNAs (average of 97 barcodes per sgRNA). We then induced Cas9 expression, measured reporter activation by immunofluorescence, and determined barcode sequences by *in situ* sequencing. All barcodes encoding targeting sgRNAs were distinguishable from control sgRNAs by HA^+^ fraction. False positive events in which HA^+^ cells were assigned control sgRNAs were used to calculate a per-cell misidentification rate of 9.4% (**Fig. 3, D and E, Methods**). We also screened the same cell library via FACS followed by deep sequencing and observed a similar enrichment of targeting sgRNAs (**Fig. 3E**), with similar representation of most barcodes in both contexts (95% within 5-fold abundance, **Fig. 3F**). We achieved comparably robust mapping of CRISPR sgRNAs to the frameshift reporter phenotype with the CROP-seq vector (4), in which the sgRNA sequence is duplicated in the 3’ UTR of an antibiotic resistance transcript and may be directly sequenced *in situ* (**Fig. S4**).

**Fig. 3:**
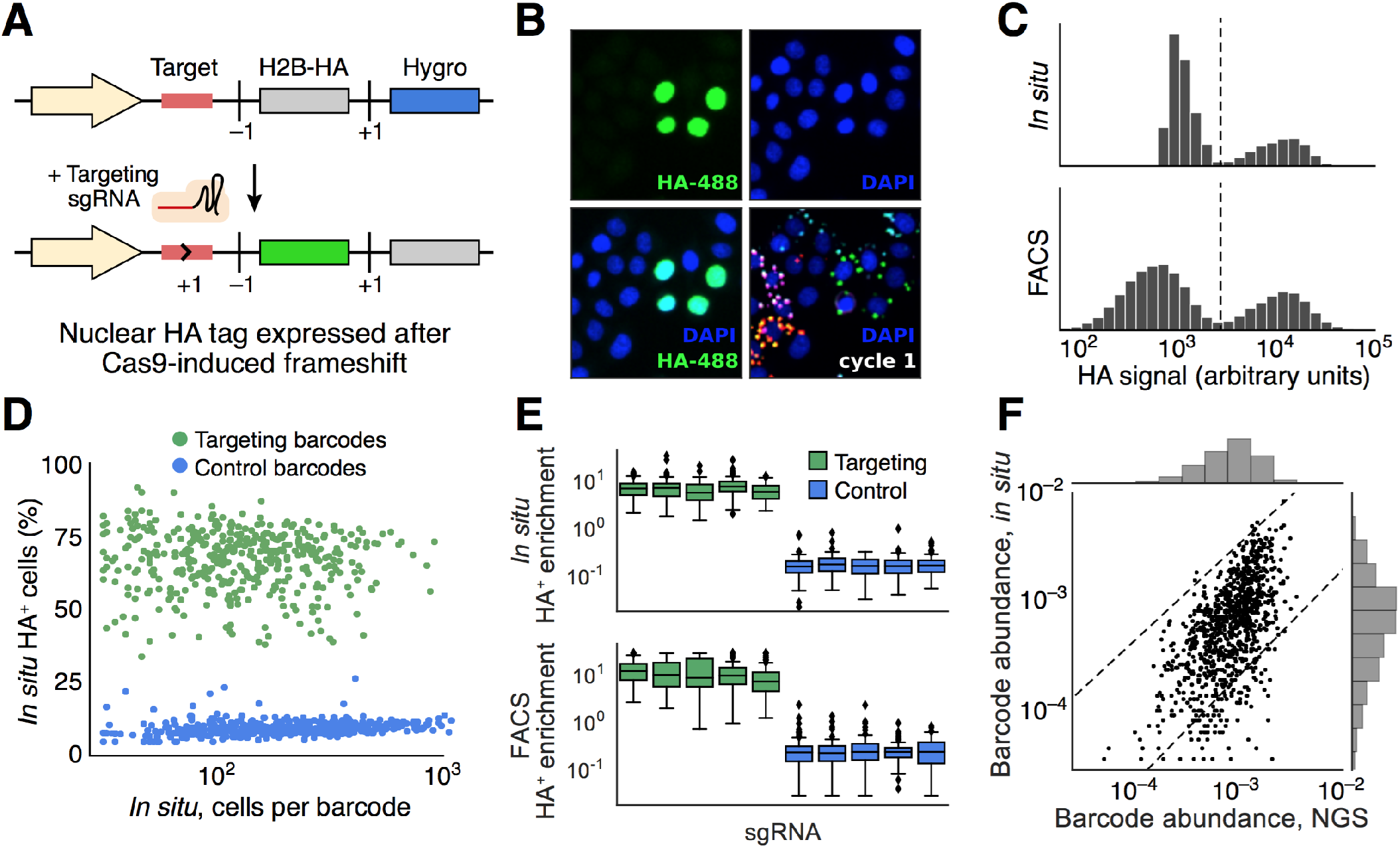
Accuracy of phenotype-to-genotype mapping assessed with a fluorescent reporter. (**A**) Schematic of a frameshift reporter that converts CRISPR-Cas9-induced indel mutations into a positive fluorescent signal. (**B**) HeLa-TetR-Cas9 cells were transduced first with the frameshift reporter and subsequently with a library of 972 barcodes, each encoding a targeting or control sgRNA. (**C**) After Cas9 induction, the frameshift reporter was activated and the resulting nuclear-localized HA epitope tag may be stained with labeled antibody and detected in each cell, either by immunofluorescence or via FACS; corresponding barcode sequences were read out by *in situ* sequencing or NGS. (**D**) Targeting and control barcodes were well separated by fraction of HA^+^ cells. (**E**) The same cell library was screened by flow sorting cells into HA^+^ and HA^-^ bins and performing next-generation sequencing of the genomically integrated barcode. (**F**) Comparison of barcode abundances measured by *in situ* sequencing or NGS (R^2^ = 0.55). The relative abundance of 95% of barcodes was within 5-fold (indicated by dashed lines).

Finally, we performed a large screen to identify genes required for activation of NF-κB, an extensively studied family of transcription factors that translocate to the nucleus in response to a host of stimuli (27, 28). We measured nuclear translocation of a p65-mNeonGreen reporter in HeLa-TetR-Cas9 cells following stimulation with either IL-1β or TNFα, cytokines that activate NF-κB via alternate pathways (**Fig. 4A**). We screened a library targeting 952 genes (3,063 sgRNAs) encompassing known NF-κB pathway-related components as well as all genes with GO annotations for ubiquitin ligase and deubiquitination activity, some of which are known to positively and negatively regulate NF-κB activation (29) (**Fig. 4A**). After stimulation with either IL-1β or TNFα, we scored the degree of p65-mNeonGreen translocation in each cell and ranked the perturbations by the deviation in their translocation score distribution from negative control sgRNAs to identify hit genes specific to IL-1β or TNFα as well as genes affecting the response to both cytokines (**Fig. 4, B and C, and Table S5, Methods**). Statistical significance was determined by permutation testing of non-targeting sgRNAs. We collected the full primary screen dataset from a single multi-well plate in which a total of 8,168,177 cells were imaged, with 3,037,909 cells retained for analysis after filtering cells based on reporter expression, nuclear morphology, and exact barcode mapping.

**Fig. 4:**
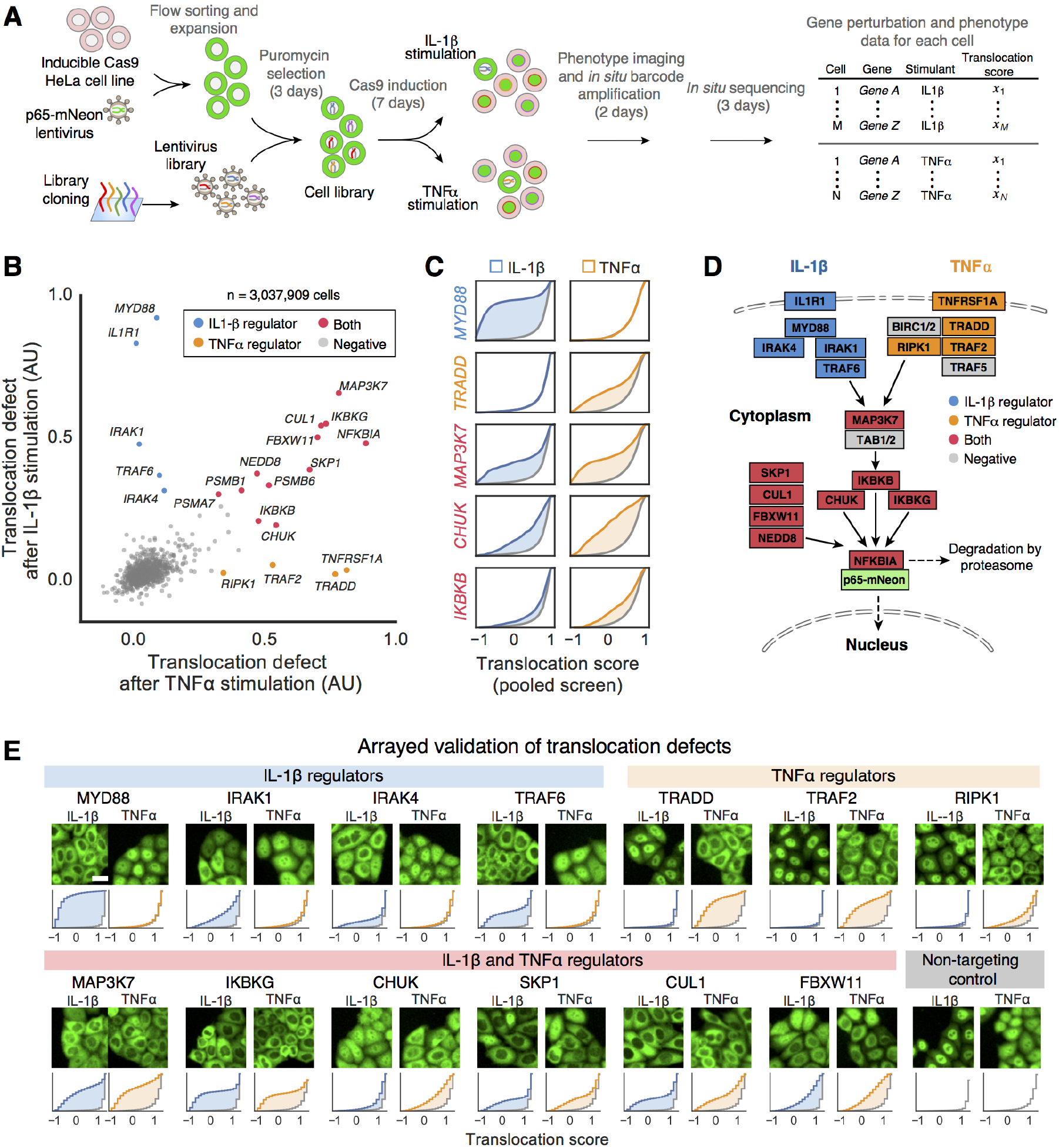
A screen for regulators of NF-κB signaling. (**A**) Workflow for CRISPR-Cas9 knockout-based screening using a fluorescently tagged reporter cell line. Screen hits were identified by the failure of p65-mNeonGreen to translocate following stimulation with IL-1β or TNFα cytokines. (**B**) Known NF-κB regulators were identified as high-ranking screen hits. Cells were assigned translocation scores based on the pixelwise correlation between mNeonGreen fluorescence and a DAPI nuclear stain. The translocation defect for a gene was defined based on the integrated difference in the distribution of translocation scores relative to non-targeting control sgRNAs across three replicate screens. (**C**) Cumulative distributions of translocation scores of known NF-κB regulators in response to both cytokines. The shaded areas depict the difference between the translocation score distributions for targeting sgRNAs and non-targeting control sgRNAs (gray). (**D**) NF-κB pathway map (KEGG HSA04064) color-coded as in (**B**). (**E**) Top-ranked genes were validated with individual CRISPR-Cas9 knockouts. Histograms show the cumulative distributions of IL-1β and TNFα-induced translocation scores for each gene knockout compared to wildtype cells (gray).

Our screen recovered known pathway components annotated by KEGG (30) for NF-κB activation by IL-1β signaling (5/5 genes), TNFα signaling (4/7 genes) and downstream components (5/7 genes), including the cytokine-specific receptors, adapter proteins, and factors that activate the shared regulator MAP3K7 (**Fig. 4D**) (28). Hits common to both cytokines included *MAP3K7* itself and its target, the IKK complex *(CHUK, IKBKB, IKBKG)*, as well as components of the SKP1-CUL1-F-box ubiquitin ligase complex and proteasome subunits, which together lead to degradation of the inhibitory IκBα protein and nuclear translocation of p65. Validation with arrayed CRISPR knockouts confirmed 17/17 top-ranked hits (**Fig. 4E and Fig. S5**), three other known pathway members *(IRAK4, RIPK1, BIRC2)*, and potentially novel hits including *BAP1, HCFC1*, and *DCUN1D1.* Phenotype strength was well correlated between the primary screening and validation data (Spearman’s ρ=0.87 (IL-1β), ρ=0.76 (TNFα)), emphasizing the quantitative nature of the primary screen (**Fig. S6**). Within the group of IL-1β-specific screening hits, *BAP1* has been previously described to deubiquitinate *HCFC1* (31), with relevance for controlling metabolism (32), ER-stress signaling (33), cell-cycle progression (34), and viral gene expression (35). However, no involvement in NF-κB signaling has been described to our knowledge. It will be interesting to further elucidate the involvement of these genes in pro-inflammatory signaling.

## Discussion

Pooled optical screens are a novel method for systematic analysis of the genetic components underpinning a broad range of spatially and temporally defined phenotypes. The workflow closely mimics conventional pooled screening and requires no specialized hardware apart from a standard automated epifluorescence microscope. Optical readout of genetic perturbations is compatible with any perturbation that can be identified by a short expressed sequence, including Cas9-mediated gene knockout, repression, and activation, as well as libraries of ORFs, protein variants, and non-coding elements barcoded with expressed tags (36, 37).

Our approach is broadly applicable across many settings. For example, multiple perturbation barcodes can be read out within the same cell (**Fig. S7**), suggesting a straightforward route to studying genetic interactions with microscopy. The potential to integrate optical screening with highdimensional morphological profiling and *in situ* multiplexed gene expression analysis (19, 38–41) raises the prospect of learning phenotypes from data rather than pre-specifying phenotypes of interest. Existing protocols for *in situ* sequencing in tissue samples (19, 41) highlight the exciting possibility of perturbing cells *in vivo* and measuring the resulting phenotypes within the native spatial context.

## Methods

### Tissue culture

HEK293FT cells (ATCC CRL-1573) were cultured in DMEM with sodium pyruvate and GlutaMAX (Life 10569044) supplemented with heat-inactivated fetal bovine serum (Seradigm 97068-085) and 100 U/mL penicillin-streptomycin (Life Technologies 15140163). All HeLa cell lines were cultured in the same media with serum substituted for 10% tetracycline-screened fetal bovine serum (Hyclone SH30070.03T).

Parental HeLa-TetR-Cas9 cells were a gift from Iain Cheeseman. In order to select an optimal clone for further experiments, single cells were sorted into a 96-well plate (Sony SH800), clonally expanded, and screened for Cas9 activity after 8 days with and without 1 μg/mL doxycycline induction. Cas9 activity was assessed by transducing each clone with pXPR_011 (Addgene #59702), a reporter vector expressing GFP and an sgRNA targeting GFP, and using FACS to read out efficiency of protein knockdown. Additionally, gene editing was directly assessed by transduced HeLa-TetR-Cas9 clones with a guide targeting TFRC. Genomic DNA was extracted from both uninduced and induced clones by resuspending in cell lysis buffer (10 mM Tris pH 7.5, 1 mM CaCl_2_, 2 mM MgCl_2_, 1 mM EDTA, and 0.2 mg/mL Proteinase K), and heating for 10 minutes at 65°C and 15 minutes at 95°C. The guide target region was amplified by PCR and sequenced on an Illumina MiniSeq. The best clones showed efficient indel generation (≥ 97%) in the presence of doxycycline and minimal cutting (≤ 2%) in its absence.

In preparation for *in situ* analysis, cells were seeded onto glass-bottom plates (6-well: Cellvis P06-1.5H-N, 24-well: Greiner Bio-one 662892, 96-well: Greiner Bio-one 665892) at a density of 50,000 cells/cm^2^ and incubated for 2 days to permit proper cell attachment, spreading, and colony formation.

### Lentiviral production

HEK293FT cells were seeded into 15-cm plates or multi-well plates at a density of 100,000 cells/cm^2^. After 20 hours, cells were transfected with pMD2.G (Addgene #12259), psPAX2 (Addgene #12260), and a lentiviral transfer plasmid (2:3:4 ratio by mass) using Lipofectamine 3000 (Thermo Fisher L3000015). Media was exchanged after 4 hours and supplemented with 2 mM caffeine 20 hours post-transfection to increase viral titer. Viral supernatant was harvested 48 hours after transfection and filtered through 0.45 μm PVDF filters (Millipore SLHVR04NL).

### Lentiviral transduction

HeLa-TetR-Cas9 cells were transduced by adding viral supernatant supplemented with polybrene (8 μg/mL) and centrifuging at 1000g for 2 hours at 33°C. At 5 hours post-infection, media was exchanged. At 24 hours post-infection, cells were passaged into media containing selection antibiotic at the following concentrations: 1 μg/mL puromycin (ThermoFisher A1113802), 300 μg/mL hygromycin (Invivogen ant-hg-1), and 300 μg/mL zeocin (ThermoFisher R25001).

For lentiviral transduction of lentiGuide-BC libraries, a carrier plasmid was utilized to minimize recombination between distant genetic elements (e.g., sgRNA and associated barcode). Libraries were packaged following the above protocol, with the library transfer plasmid diluted in integration-deficient carrier vector pR_LG (1:10 mass ratio of library to carrier, Addgene #112895) *(25)*.

### Library cell line validation

For library transductions, multiplicity of infection was estimated by counting colonies after sparse plating and antibiotic selection. Genomic DNA was also extracted for NGS validation of library representation.

### Next generation sequencing of libraries

Genomic DNA was extracted using an extraction mix as described above. Barcodes and sgRNAs were amplified by PCR from a minimum of 1e6 genomic equivalents per library using NEBNext 2X Master Mix (initial denaturation 5 minutes at 98°C, followed by 28 cycles of annealing for 10 seconds at 65°C, extension for 25 seconds at 72°C, and denaturation for 20 seconds at 98°C).

### Library design and cloning

A set of 12-nt barcodes was designed by selecting 83,314 barcodes from the set of 16.7 million possible 12-nt sequences by filtering for GC content between 25% and 75%, no more than 4 consecutive repeated bases, and minimum substitution and insertion/deletion edit distance (Levenshtein distance) of 3 between any pair of barcodes. Ensuring a minimum edit distance is useful for detecting and correcting errors, which arise mainly from DNA synthesis and *in situ* reads with low signal-to-background ratios. The E-CRISP web tool was used to select sgRNA sequences targeting genes of interest. Barcode-sgRNA pairs were randomly assigned and co-synthesized on a 125-nt 90K oligo array (CustomArray) (**table S4**). Individual libraries were amplified from the oligo pool by dial-out PCR (42) and cloned into lentiGuide-BC or lentiGuide-BC-CMV (the latter contains the CMV promoter instead of the EF1a promoter) via two steps of Golden Gate assembly using BsmBI and BbsI restriction sites. Libraries were transformed in electrocompetent cells (Lucigen Endura) and grown in liquid culture for 18 hours at 30°C before extracting plasmid DNA. The sgRNA-barcode association was validated by Sanger sequencing individual colonies from the final library.

### Padlock-based RNA detection

Preparation of targeted RNA amplicons for *in situ* sequencing was adapted from published protocols with modifications to improve molecular detection efficiency and amplification yield (19, 43). Cells were fixed with 4% paraformaldehyde (Electron Microscopy Sciences 15714) for 30 minutes, washed with PBS, and permeabilized with 70% ethanol for 30 minutes. Permeabilization solution was carefully exchanged with PBS-T wash buffer (PBS + 0.05% Tween-20) to minimize sample dehydration. Reverse transcription mix (1x RevertAid RT buffer, 250 μM dNTPs, 0.2 mg/mL BSA, 1 μM RT primer, 0.8 U/μL Ribolock RNase inhibitor, and 4.8 U/μL RevertAid H minus reverse transcriptase) was added to the sample and incubated for 16 hours at 37°C. Following reverse transcription, cells were washed 5 times with PBS-T and post-fixed with 3% paraformaldehyde and 0.1% glutaraldehyde for 30 minutes at room temperature, then washed with PBS-T 5 times. After this step, cells expressing p65-mNeonGreen were imaged. Samples were thoroughly washed again with PBS-T, incubated in a padlock probe and extension-ligation reaction mix (1x Ampligase buffer, 0.4 U/μL RNase H, 0.2 mg/mL BSA, 10 nM padlock probe, 0.002 U/μL TaqIT polymerase, 0. 5 U/μL Ampligase and 50 nM dNTPs) for 5 minutes at 37°C and 90 minutes at 45°C, and then washed 2 times with PBS-T. Circularized padlocks were amplified with rolling circle amplification mix (1x Phi29 buffer, 250 μM dNTPs, 0.2 mg/mL BSA, 5% glycerol, and 1 U/μL Phi29 DNA polymerase) at 30°C for either 3 hours or overnight.

### *In situ* sequencing

Rolling circle amplicons were prepared for sequencing by hybridizing a mix containing sequencing primer pSBS_lentiGuide-BC or pSBS_CROPseq (1 μM primer in 2X SSC + 10% formamide) for 30 minutes at room temperature. Barcodes were read out using sequencing-by-synthesis reagents from the Illumina MiSeq 500 cycle Nano kit (Illumina MS-103-1003). First, samples were washed with incorporation buffer (Nano kit PR2) and incubated for 3 minutes in incorporation mix (Nano kit reagent 1) at 60°C. Samples were then thoroughly washed with PR2 at 60°C (6 washes for 3 minutes each) and placed in 200 ng/mL DAPI in 2x SSC for fluorescence imaging. Following each imaging cycle, dye terminators were removed by incubation for 6 minutes in Illumina cleavage mix (Nano kit reagent 4) at 60°C, and samples were thoroughly washed with PR2.

### Fluorescence microscopy

All images were acquired using a Ti-E Eclipse inverted epifluorescence microscope (Nikon) with automated XYZ stage control and hardware autofocus. An LED light engine (Lumencor Sola SE FISH II) was used for fluorescence illumination and all hardware was controlled using Micromanager software (44). *In situ* sequencing cycles were imaged using a 10X 0.45 NA CFI Plan Apo Lambda objective (Nikon) with the following filters (Semrock) and exposure times for each base: G (excitation 534/20 nm, emission 572/28 nm, dichroic 552 nm, 200 ms); T (excitation 575/25 nm, emission 6158/24 nm, dichroic 596 nm, 200 ms); A (excitation 635/18 nm, emission 680/42 nm, dichroic 652 nm, 200 ms); C (excitation 661/20 nm, emission 732/68 nm, dichroic 695 nm, 800 ms).

### Image analysis

Images of cell phenotype and *in situ* sequencing of perturbations were aligned using cross-correlation of DAPI-stained nuclei. Nuclei were detected using local thresholding and watershed-based segmentation. Cells were typically segmented using local thresholding of cytoplasmic background and assignment of pixels to the nearest nucleus by the fast-marching method. Frameshift reporter and NF-κB translocation phenotypes were quantified by calculating pixel-wise correlations between the nuclear DAPI channel and 488 channel (HA stain or p65-mNeonGreen, respectively).

Sequencing reads were detected by applying a Laplacian-of-Gaussian linear filter (kernel width σ = 1 pixel), calculating the per-pixel standard deviation over sequencing cycles, averaging over color channels, and finding local maxima. The base intensity at each cycle was defined as the maximum value in a 3×3 pixel window centered on the read. A linear transformation was then applied to correct for optical crosstalk and intensity differences between color channels. Finally, each base was called according to the channel with maximum corrected intensity, and a per-base quality score was defined as the ratio of intensity for the maximum channel to total intensity for all channels. The output of the sequencing image analysis was a FASTQ file recording each sequencing read along with the identity of the overlapping cell, quality score per base, and spatial location.

Additional details and source code available on Github at https://github.com/blaineylab/OpticalPooledScreens.

### Frameshift reporter screen

HeLa-TetR-Cas9 cells were stably transduced at MOI >2 with pL_FR_Hygro and selected with hygromycin for 7 days to generate the HeLa-TetR-Cas9-FR cell line. Cells transduced with the pL_FR_Hygro lentiviral vector express an open reading frame consisting of a 50-nt frameshift reporter target sequence, followed by an H2B histone coding sequence with C-terminus HA epitope tag (+1 frameshift), followed by a second H2B sequence with C-terminus myc tag (+0 frameshift) and hygro antibiotic resistance cassette (+0 frameshift). The H2B-HA, H2B-myc, and hygromycin resistance sequences are preceded by self-cleaving 2A peptides in the same reading frame. Before generation of Cas9-mediated indel mutations, cells express the coding sequences with +0 frameshift. Subsequent activation of the reporter by co-expression of Cas9 and a targeting sgRNA leads to mutations in the target sequence, which may alter the downstream reading frame. A frameshift of +1 leads to expression of the H2B-HA protein, which can be visualized by immunofluorescence and detected by microscopy or flow cytometry. Integration of multiple copies of reporter per cell increases the likelihood of generating a +1 frameshift in at least one copy.

HeLa-TetR-Cas9-FR cells were used to screen targeting and control sgRNAs. A barcoded sgRNA perturbation library with 972 barcodes, each encoding one of 5 targeting or 5 control sgRNAs, was synthesized and cloned into lentiGuide-BC-CMV. This library was transduced into HeLa-TetR-Cas9-FR cells at MOI < 0.05 in three replicates, which were independently cultured and screened. Following 4 days of puromycin selection, cells were collected to validate library representation by NGS. Cas9 expression was induced by supplementing the culture media with 1 μg/mL doxycycline for 6 days. Cells were then split for screening either via *in situ* sequencing or by FACS.

For *in situ* screening, 500,000 cells were seeded into each well of a glass-bottom 6-well plate (CellVis). After two days of culture, *in situ* padlock detection and sequencing were carried out as above, with the modification that prior to sequencing-by-synthesis, cells were immunostained to detect frameshift reporter activation by blocking and permeabilizing with 3% BSA + 0.5 % Triton X-100 for 5 minutes, incubating in rabbit anti-HA (1:1000 dilution in 3% BSA) for 30 minutes, washing with PBS-T and incubating with goat anti-rabbit F(ab’)2 fragment Alexa 488 (CST 4412S, 1:1000 dilution in 3% BSA) for 30 minutes. Samples were changed into imaging buffer (200 ng/mL DAPI in 2X SSC), and phenotype images were acquired.

FACS screening was carried out by fixing cells with 4% PFA, permeabilizing with 70% ice-cold ethanol, and immunostaining with the same anti-HA primary and secondary antibodies and dilutions used for *in situ* analysis. Cells were sorted into HA^+^ and HA^-^ populations (Sony SH800) and genomically integrated perturbations were sequenced as described above. The enrichment for each barcoded perturbation was defined as the ratio of normalized read counts.

To estimate the rate at which cells are assigned an incorrect perturbation barcode, we first assumed all HA^+^ cells mapped to a non-targeting control sgRNA (4.7%) were false positive events due to incorrect barcode assignment (supported by the very low false positive rate (<0.001%) of the frameshift reporter itself, measured for a single perturbation in arrayed format). However, as incorrect barcode assignments were equally likely to map an HA^+^ cell to a targeting or control sgRNA, we estimated the misidentification rate to be twice as large, or 9.4%.

### NF-κB pooled screen

HeLa-TetR-Cas9 cells were transduced with pR14_p65-mNeonGreen, a C-terminal fusion of p65 with a bright monomeric green fluorescent protein (Allele Biotechnology). Fluorescent cells were sorted by FACS (Sony SH800) and resorted to select for cells with stable expression. This reporter cell line was further transduced with a 4,063-barcode sgRNA library (962 genes targeted (952 detected), 866 barcodes assigned to non-targeting controls) in lentiGuide-BC-CMV. Cells were selected with puromycin for 4 days and library representation was validated by NGS.

Cas9 expression was induced with 1 μg/mL doxycycline and cells were seeded onto 6-well glass-bottom plates 2 days prior to translocation experiments. The total time between Cas9 induction and performing the NF-κB activation assay was 7 days. Cells were stimulated with either 30 ng/mL TNFα or 30 ng/mL IL-1β for 40 minutes prior to fixation with 4% paraformaldehyde for 30 minutes and initiation of the *in situ* sequencing protocol. Translocation phenotypes were recorded after the post-fixation step by exchanging for imaging buffer (2X SSC + 200 ng/mL DAPI) and imaging the nuclear DAPI stain and p65-mNeonGreen. After phenotyping, the remainder of the *in situ* sequencing protocol (gap-fill and rolling circle amplification) and 12 bases of sequencing-by-synthesis were completed.

Nuclei of individual cells were segmented by thresholding background-subtracted DAPI signal and separating the resulting regions using the watershed method. Cells with at least one read exactly matching a library barcode were retained for analysis. In order to remove mitotic cells and cells with abnormally high or low reporter expression, cells were further filtered based on nuclear area, maximum DAPI signal, and mean p65-mNeonGreen signal. Pixel-wise DAPI-mNeonGreen correlation within the segmented nuclear region, described above, was used to define the translocation score for each cell as it most clearly separated perturbations against known NF-κB genes from non-targeting controls. The phenotypic effects of perturbations targeting known NF-κB genes ranged from a large increase in fully untranslocated cells (e.g., *MAP3K7*) to more subtle negative shifts in the distribution of scores (e.g., *IKBKB*).

To capture a broad range of effect size, we calculated an sgRNA translocation defect in a given replicate by computing the difference in translocation score distribution compared to non-targeting controls (the shaded area in **Fig. 4, C and E**). We found this metric performed better at separating known genes from controls than the often-used Kolmogorov-Smirnov distance. We defined the gene translocation defect as the second-largest sgRNA translocation defect for sgRNAs targeting that gene. This statistic helps reduce the false positive rate due to clonal effects (integration of an sgRNA into a cell that is defective in translocation) which are independent among sgRNAs and screen replicates, as well as false negatives due to inefficient sgRNAs.

A permutation test was used to calculate p-values for the gene translocation defects. Random subsets of sgRNA translocation defects were sampled from non-targeting controls to build a null distribution (3 sgRNAs per replicate, repeated 100,000 times). The cumulative null distribution was used to determine p-values for the gene translocation defects. Hits at an estimated FDR <10% and <20% were identified using the Benjamini-Hochberg procedure (**Table S5**). KEGG-annotated genes were defined as members of KEGG pathway HSA04064 (NF-kappa B signaling pathway) between IL-1β or TNFα and p65/p60.

## Acknowledgments

We thank Rhiannon Macrae, Iain Cheeseman, Aviv Regev, Fei Chen, Arjun Raj, Nir Hacohen, Sara Jones, Emily Botelho, and members of the Blainey and Zhang labs for critical feedback and discussions. We thank Leslie Gaffney for assistance with figures. We gratefully acknowledge Linyi Gao for the gift of the integration-deficient copackaging plasmid pR_LG.

## Author contributions

D.F. designed the approach with input from all authors. D.F., A. S., A.M., A.J.G., R.J.C. and J.S.B performed experiments. D. F. analyzed data. J.S.B and D.F. designed the NF-κB screen. P.C.B. and F.Z. supervised the research. D.F., A.S., and P.C.B. wrote the manuscript with contributions from all authors.

## Online software and protocols

Python scripts, including a complete pipeline from example data to sequencing reads, will be made available on Github, along with plasmid sequences and updated protocols at https://github.com/blaineylab/OpticalPooledScreens

---

**Fig. S1:**
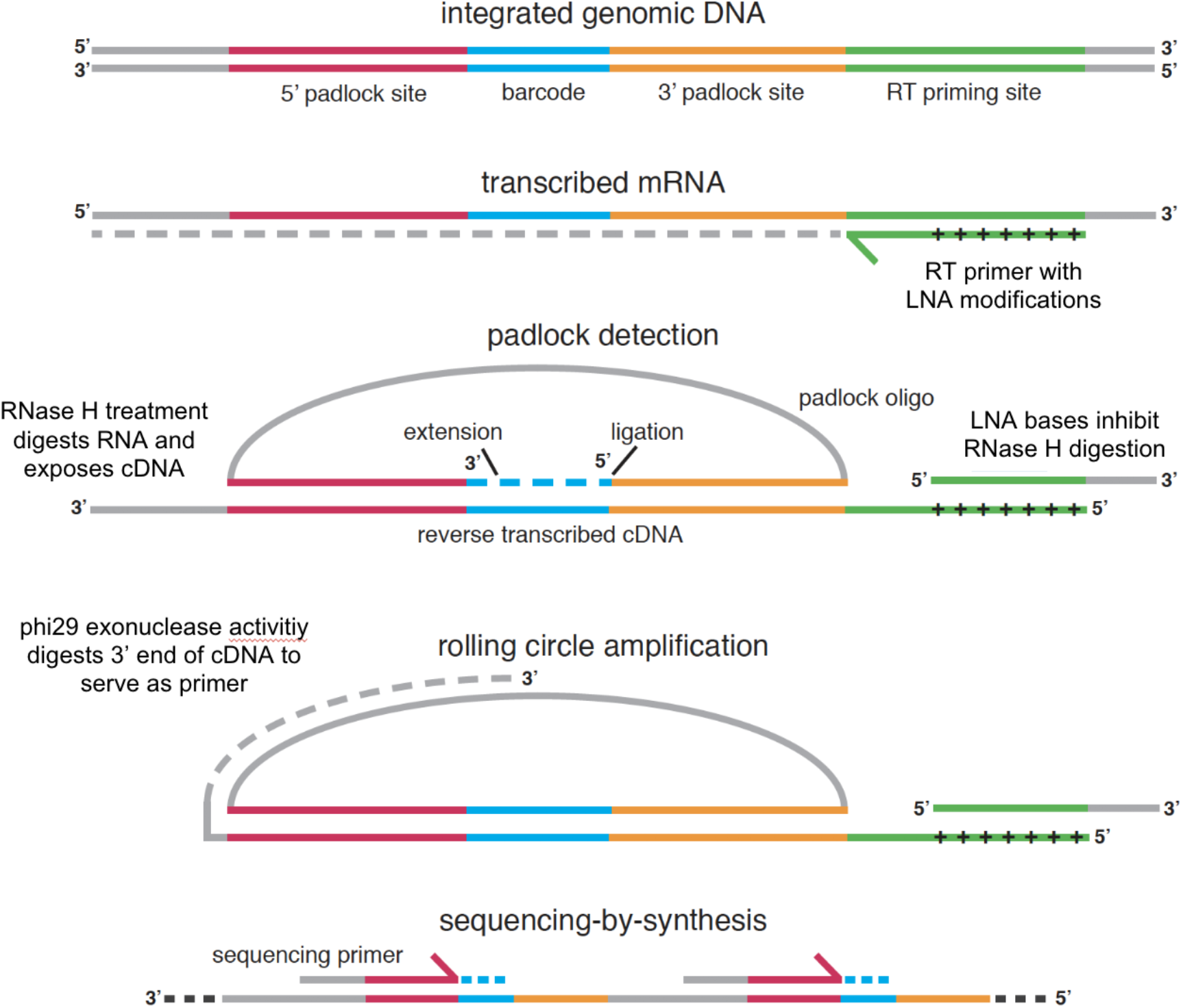
Workflow for padlock detection and sequencing-by-synthesis. In order to determine the identity of the lentiviral vector integrated in each cell, all cellular RNAs are first fixed in place by formaldehyde treatment. A reverse transcription primer containing locked nucleic acid (LNA) bases is hybridized to the mRNA containing the barcode sequence. Complementary DNA (cDNA) is generated using a reverse transcriptase lacking RNase activity, producing an RNA-DNA hybrid. The cells are then fixed once again (“post-fixed”) with a mixture of formaldehyde and glutaraldehyde to improve cDNA retention. A single reaction mix containing RNase H, a DNA polymerase lacking strand displacement activity, a DNA ligase, and a padlock DNA oligonucleotide is then added. Digestion of the RNA strand exposes the cDNA bases, allowing the padlock to hybridize to the cDNA at sites flanking the barcode. The DNA polymerase extends the padlock, copying the barcode sequence, but does not strand displace the 5’ annealed padlock arm. Once extended, the padlock is then ligated into a single-stranded DNA circle. During this step, the cDNA is retained in place via hybridization to the RNA strand at the LNA-modified bases within the RT primer, which inhibit RNase H digestion. Phi29 polymerase is used to perform rolling circle amplification of the circularized padlock. The 3’ exonuclease activity of Phi29 polymerase digests the single-stranded portion of the cDNA strand, generating a primer for rolling circle amplification. The amplified single-stranded DNA product contains tandem repeats of the padlock adaptor sequences and barcode, which can be read out by sequencing-by-synthesis. The overall protocol provides a high level of sequence specificity, conferred by hybridization of the RT primer to a unique priming site, hybridization of the padlock to the flanking sites, the preference of the ligase to act only on exactly matched DNA, and sequencing-by-synthesis of the cell-derived barcode sequence itself.

**Fig. S2:**
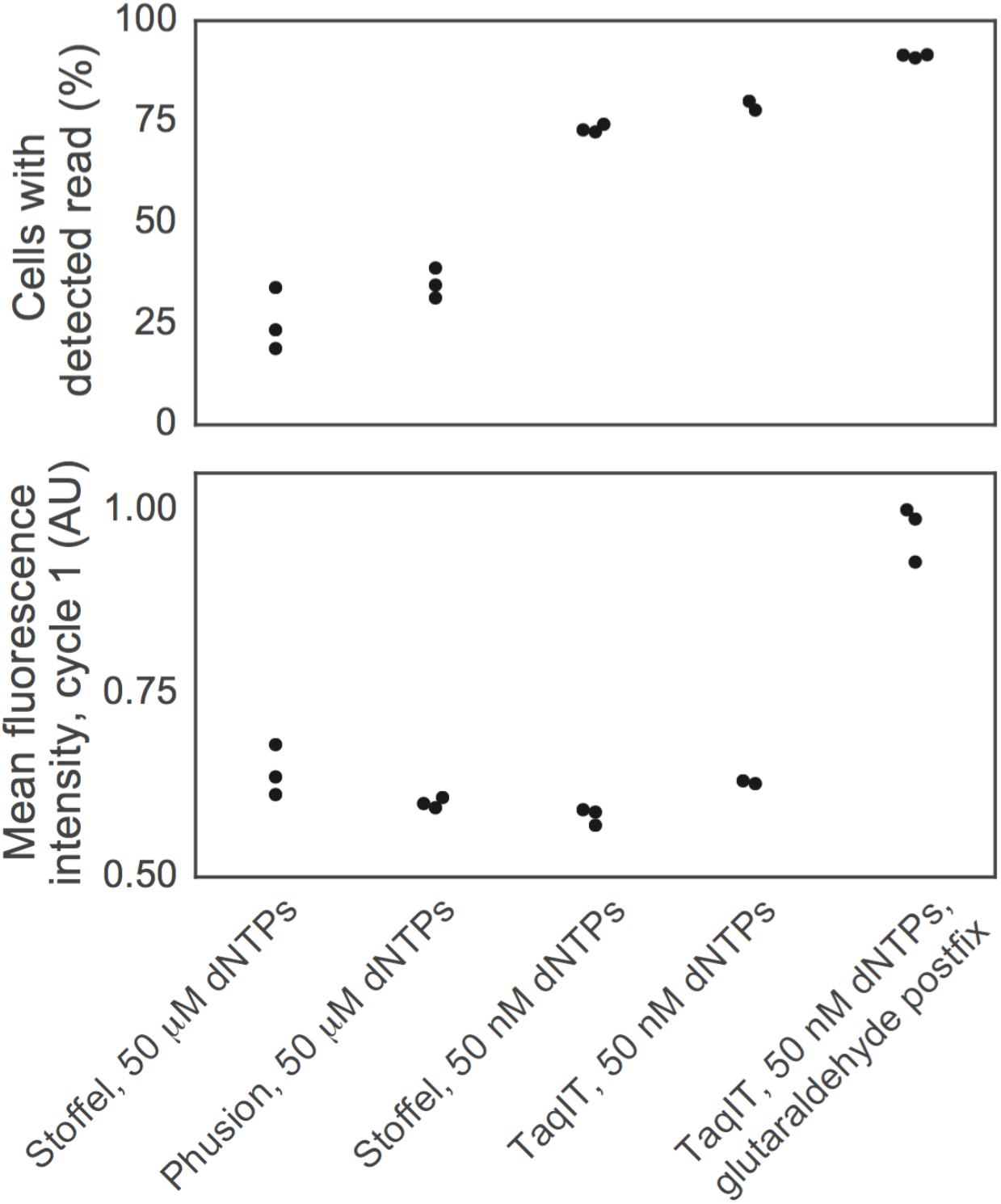
Optimization of padlock detection efficiency and amplification yield. Padlock detection efficiency was increased more than two-fold compared to literature protocols (*19, 45*) by optimizing the dNTP concentration and polymerase used for the padlock extension-ligation reaction. A striking improvement in detection efficiency was observed when using Stoffel fragment with a dNTP concentration 1000-fold less than previously published (19). Although Stoffel fragment has been discontinued by its manufacturer, we obtained similar results with another commercially available truncation mutant of *Taq* polymerase (Qiagen TaqIT). While optimizing post-fixation conditions for detection efficiency, we observed that modifying the standard 4% formaldehyde fixative to 3% formaldehyde and 0.1% glutaraldehyde led to a dramatic increase in the yield of overall fluorescence signal from each detected barcode. We presume the improvement was due to an increase in the efficiency of rolling circle amplification, although no specific mechanism was identified. The protocol comparison was performed on a single multi-well plate, using HeLa-TetR-Cas9 cells transduced with lentiGuide-BC. Each data point represents a technical replicate of the *in situ* protocol.

**Fig. S3:**
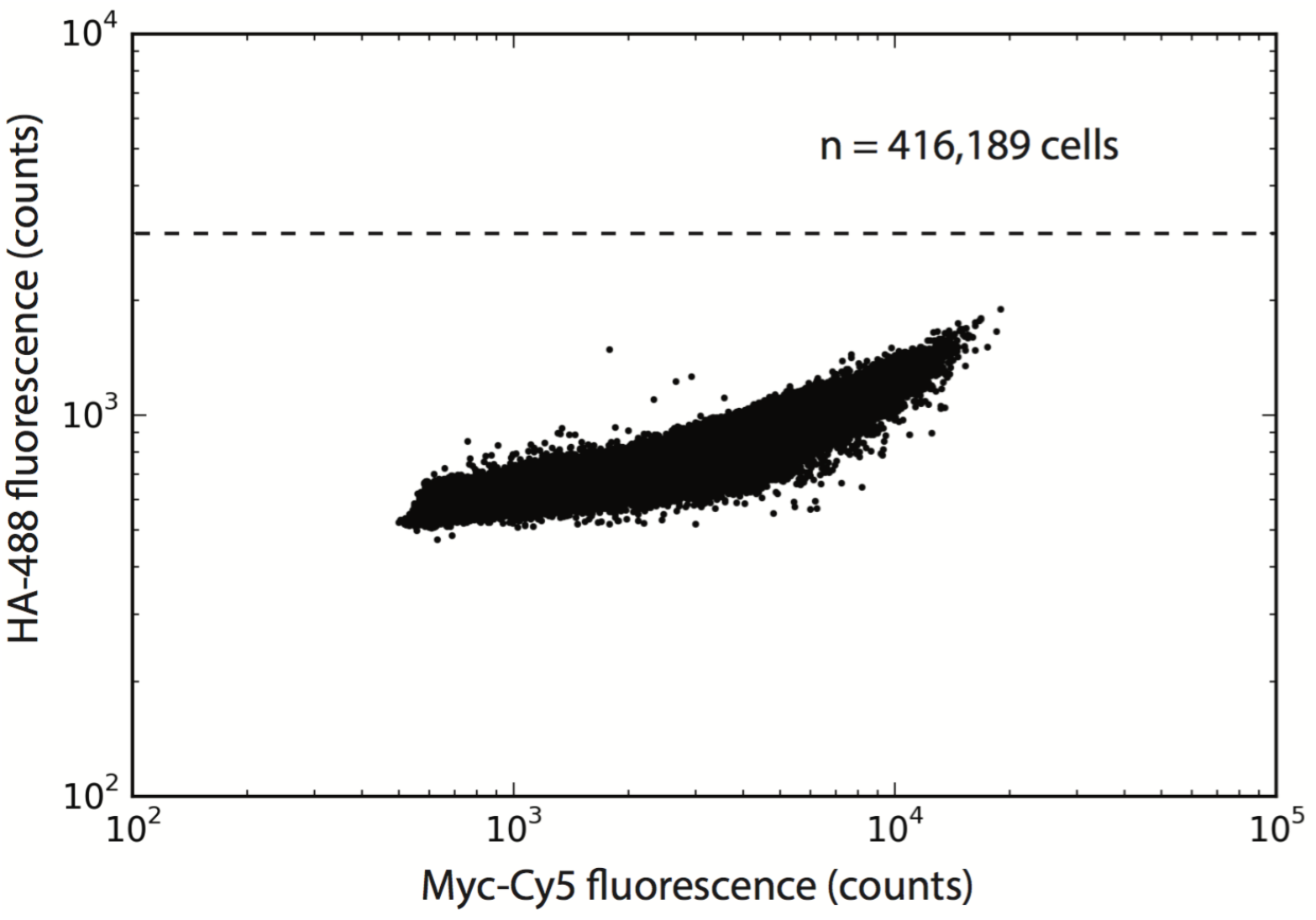
Frameshift reporter background. The frameshift reporter was read out by microscopy in HeLa-TetR-Cas9-FR cells in the absence of a targeting sgRNA. A *myc* epitope tag in the original, unedited frame was stained to confirm expression levels. The reporter was found to have a very low background, with zero false positives observed among >400,000 cells (dashed line indicates threshold for defining HA^+^ cells).

**Fig. S4:**
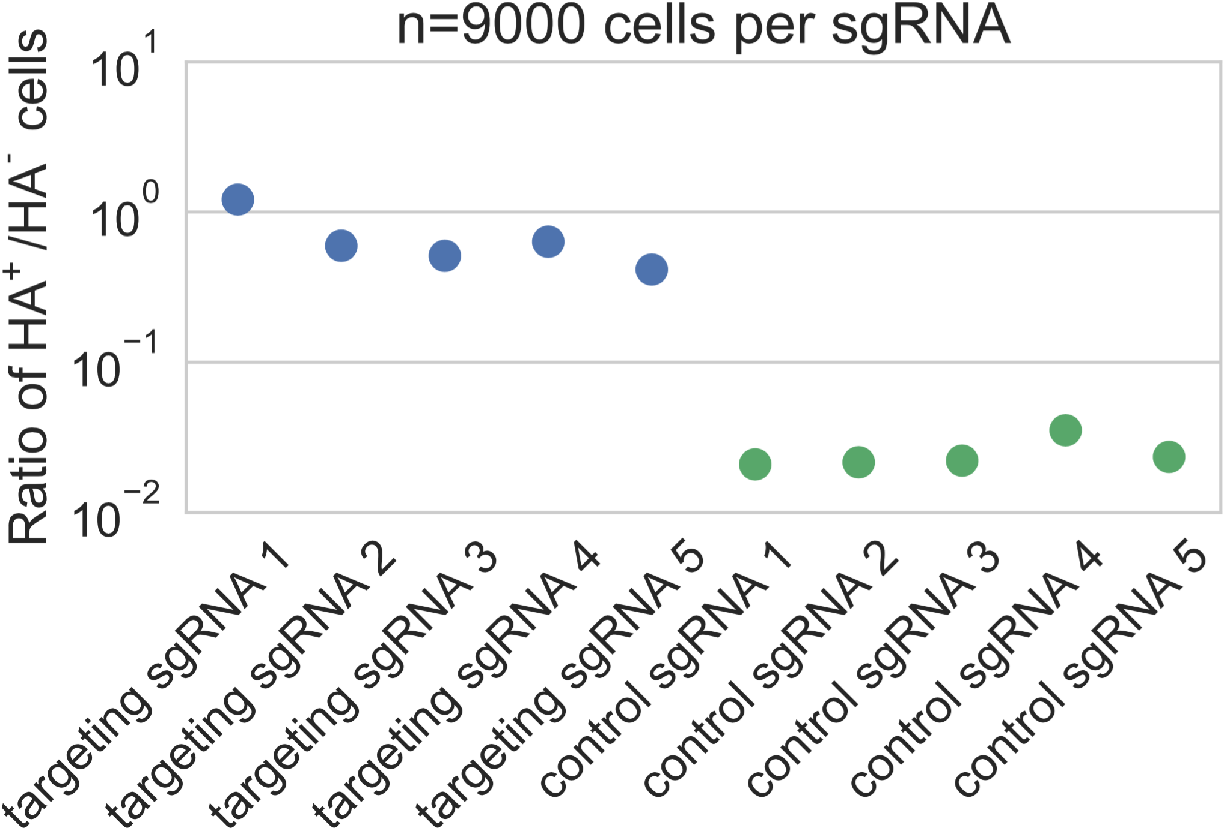
Validation of CROPseq-Puro vector in frameshift reporter cells. A pool of 5 targeting and 5 control sgRNAs were inserted into CROPseq-Puro and transduced into HeLa-Cas9 frameshift reporter cells at MOI ~10%. After Cas9 induction and frameshift reporter activation, cells were scored for nuclear-localized HA signal. The sgRNA sequence duplicated in the antibiotic resistance cassette was directly sequenced *in situ* over 4 cycles. The ratio of HA^+^ cells (19.4% of all cells) to HA^-^ cells is shown. The per-read mapping rate was 94% (exact matches to 10/256 possible 4-nt sgRNA prefixes).

**Fig. S5:**
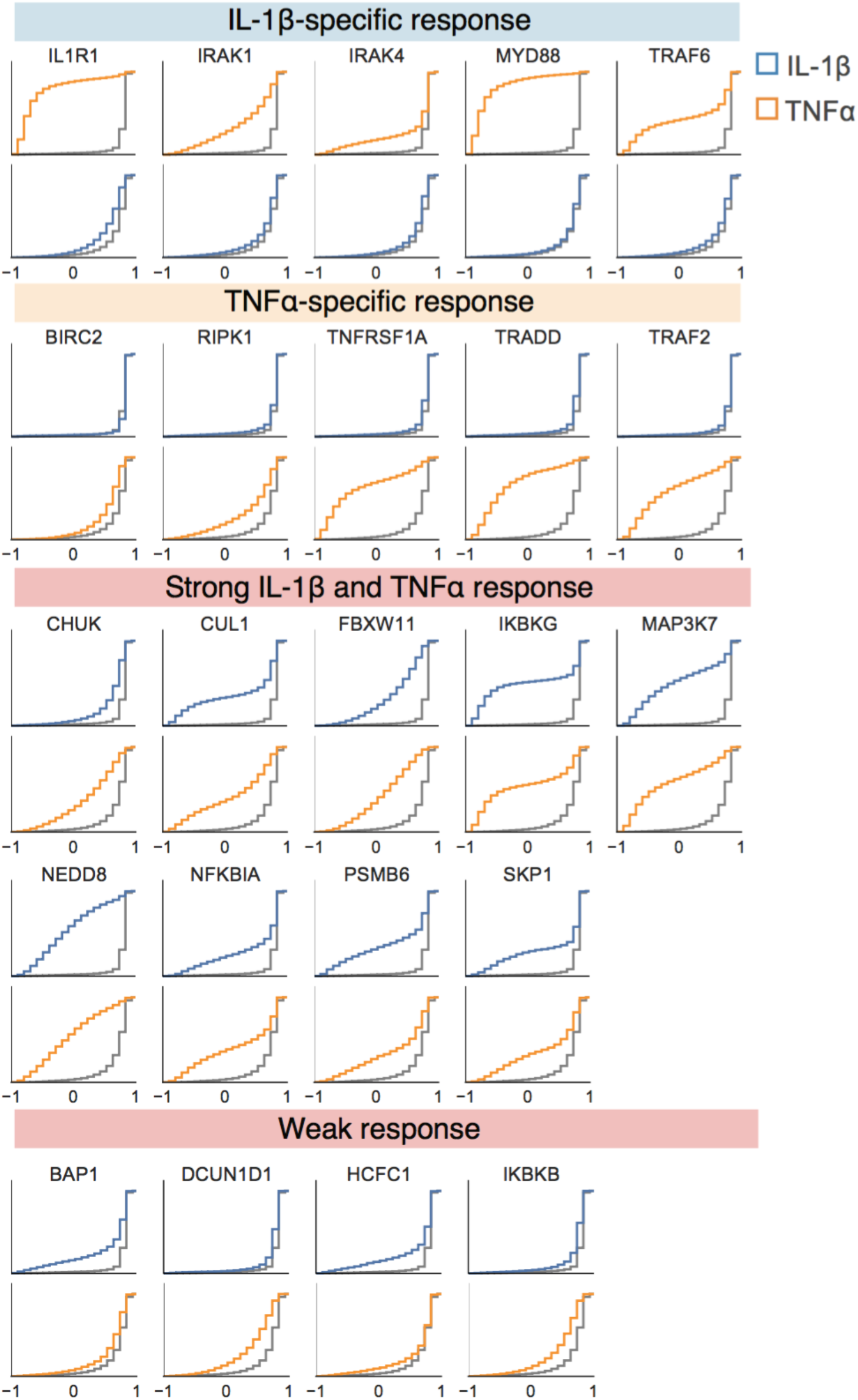
Arrayed validation of NF-κB screen hits. Individual CRISPR knockout gene perturbations were cloned and transduced into HeLa-TetR-Cas9-p65-mNeonGreen cells for validation of primary screen hits. Induction of Cas9, stimulation by IL-1β and TNFα, and translocation score analysis were performed as in the primary screen. Histograms depict the distribution of translocation scores for each knockout.

**Fig. S6:**
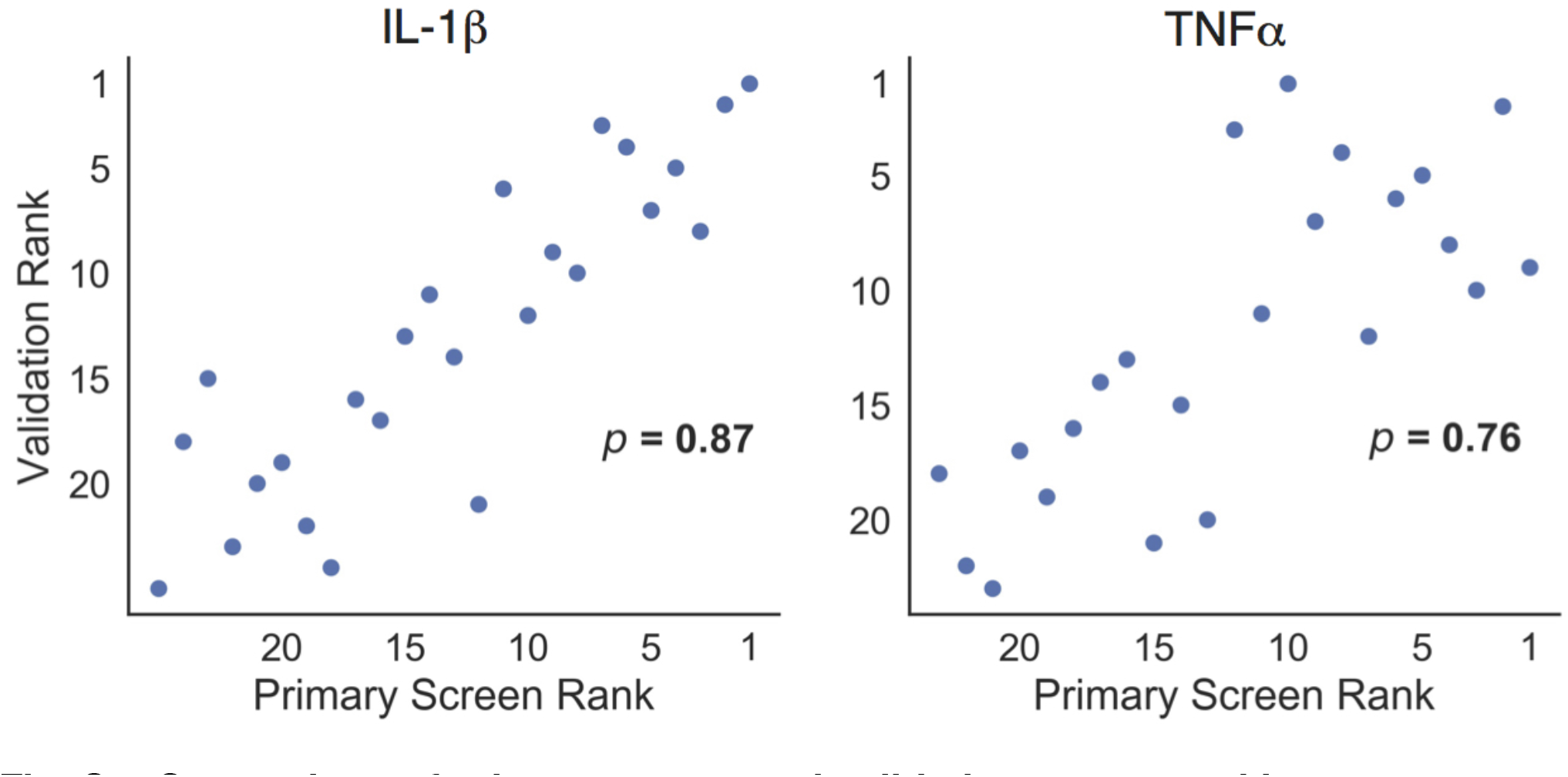
Comparison of primary screen and validation screen rankings. For both IL-1β and TNFα, primary screen gene rankings correlate well (Spearman’s p > 0.75) with rankings in validation screen of single-gene CRISPR-Cas9 knockouts. Proteasome subunits are not shown as they exhibited severe negative fitness effects in arrayed validation experiments, likely biasing the surviving cells to those with incomplete protein knockout.

**Fig. S7:**
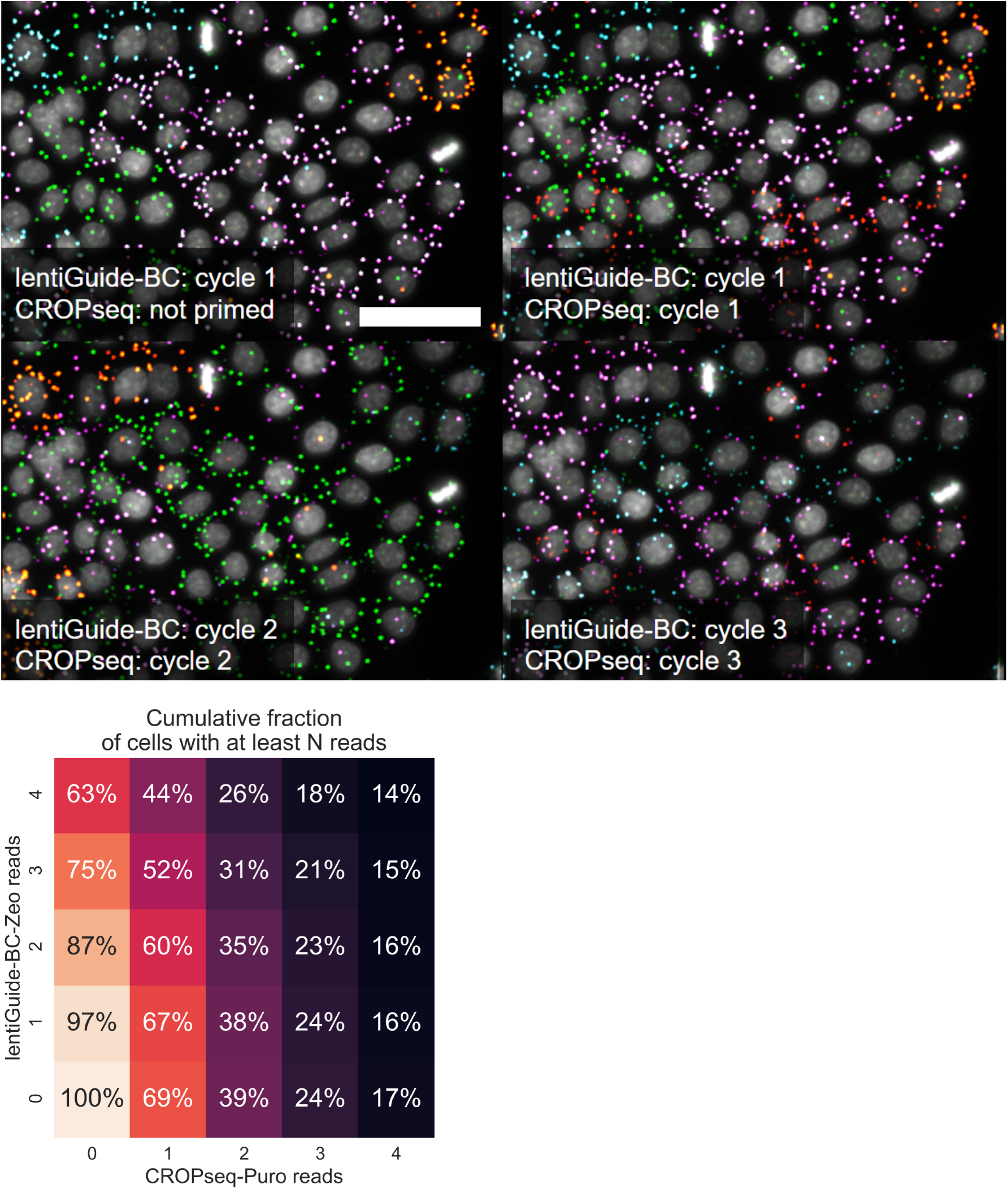
Detection of combinatorial perturbations. Multiple perturbations can be delivered via separate lentiviral vectors and detected in the same cell. HeLa-TetR-Cas9 cells were sequentially transduced with lentiGuide-BC (containing a pool of 40 barcodes) and CROPseq-Puro (containing a pool of 10 sgRNAs). The *in situ* padlock detection protocol was the same as for individual vectors, except the RT primers and padlock probes targeting each vector were mixed at equal ratios. Images were acquired at 20X magnification to improve separation of multiple barcode spots per cell (scale bar = 50 μm). The first cycle of *in situ* sequencing used only a sequencing primer targeting lentiGuide-BC. Subsequent cycles used sequencing primers targeting both constructs (top). The cumulative fraction of cells with at least N reads from each library can be calculated (bottom). For example, a total of 35% of all cells imaged had 2 or more reads from each library (heatmap position 2,2).

**Table S1:**
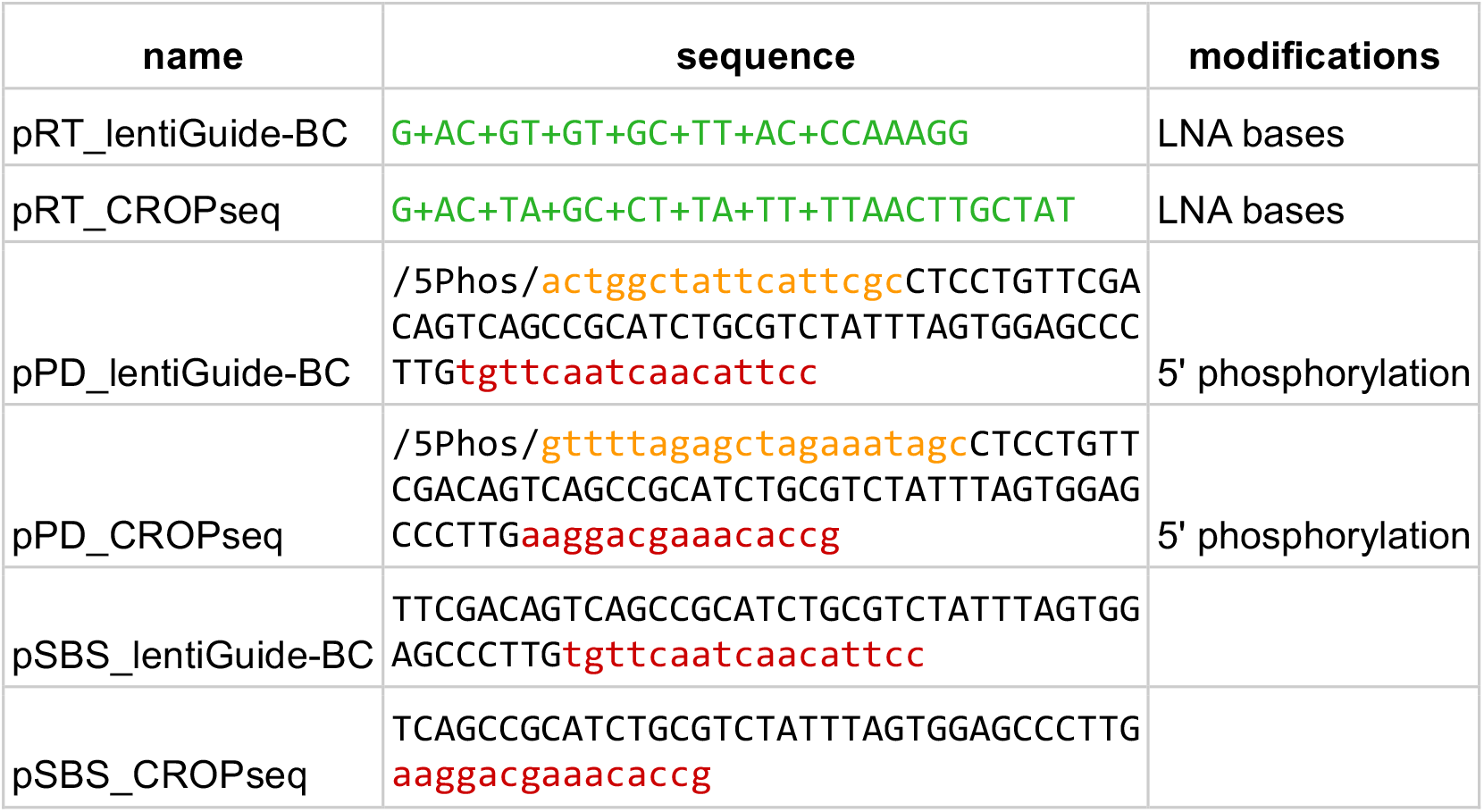
Oligo sequences used for padlock detection and sequencing-by-synthesis. Sequences of reverse transcription primer (pRT), padlock probe (pPD), and sequencing-by-synthesis primer (pSBS) used for detecting barcodes in cells transduced with the lentiGuide-BC or CROP-seq vectors. Note that LNA-modified bases in the reverse transcription primer are essential to retain the linkage between the cDNA and fixed RNA, preventing diffusion of cDNA out of the cell (19, 43).

**Table S2:** Screening throughput. The screening approach described in this study uses fully pooled protocols for library construction, cloning, and cell culture. As a result, practical limitations to screen scale come mainly from the time required to image large numbers of cells and the cost of reagents for padlock detection and *in situ* sequencing. Supplementary Table 2 summarizes the scale and throughput of the NF-κB screen performed, as well as two hypothetical screens using either low-density cell culture or a genome-scale perturbation library. The key figure determining both throughput and cost is the total surface area processed, which affects the volume of reagents used and the time per sequencing cycle. The NF-κB screen was read out from a total of ~8 million cells in a single 6-well plate.

|  | perturbations<br>(e.g., sgRNAs) | screen<br>replicates | cell<br>coverage | mapped<br>cells | cells<br>(total) | cell density<br>(cells / cm<sup>2</sup>) | sequencing<br>cycle (hours) | # of<br>cycles | total readout<br>time (days) | reagent cost<br>(direct) |
| --- | --- | --- | --- | --- | --- | --- | --- | --- | --- | --- |
| NF-κB screen in this study<br>(3 replicates each for IL1b and TNFa) | 4,000 | 6 | 130 | 3,120,000 | 8,320,000 | 175,000 | 6 | 12 | 3 | \$1,485 |
| 4,000 perturbation screen with<br>low-density cells | 4,000 | 3 | 150 | 1,800,000 | 4,800,000 | 25,000 | 21 | 12 | 11 | \$5,997 |
| Genome-scale CRISPR screen with<br>3 replicates | 60,000 | 3 | 200 | 36,000,000 | 96,000,000 | 175,000 | 59 | 12 | 30 | \$17,134 |

**Table S3:**
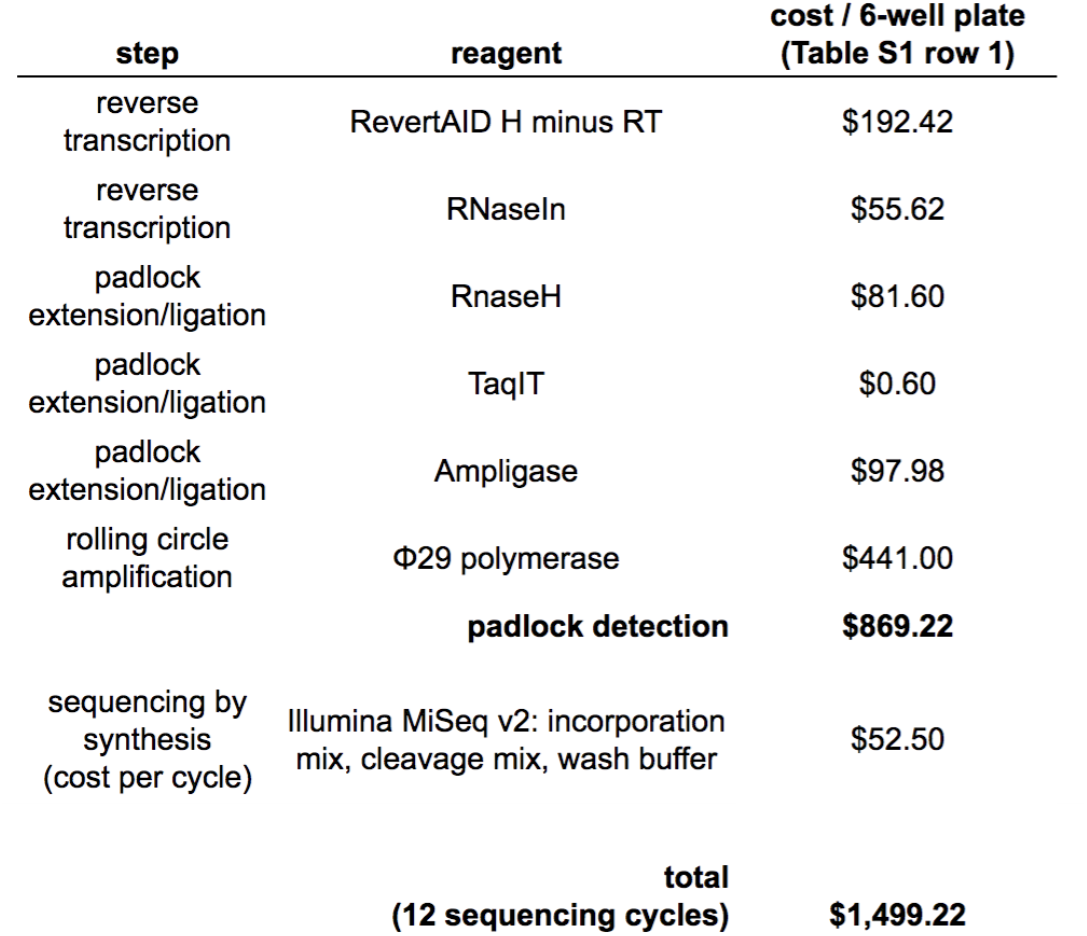
Reagent cost estimate for *in situ* barcode readout. The major reagent costs (direct) are shown. The cost per 6-well plate corresponds to the NF-κB screen in Fig 4 and the first row of Supplementary Table 2.

## Table S4: (separate file)

Libraries of paired barcodes and sgRNAs were amplified from oligo pools and cloned into lentiGuide-BC or lentiGuide-BC-CMV (Methods).

## Table S5: (separate file)

Each gene was assigned a score for lack of p65-mNeonGreen translocation in response to IL-1 β and TNFα stimulation. Gene p-values were calculated by repeatedly sampling sets of three permuted non-targeting sgRNAs to generate gene-level null translocation scores. Genes scored as hits at estimated FDR <10% or <20% were identified by the Benjamini-Hochberg procedure (Methods).

## Movie S1

Example data showing a complete 10X magnification field of view with 12 cycles of sequencing, taken from the same dataset used in Fig. 2.

